# A novel molecular sexing method for Greater One-horned Rhinoceros (*Rhinoceros unicornis*) for reliable demographic assessments

**DOI:** 10.1101/2025.05.21.655307

**Authors:** Shrewshree Kumar, Tista Ghosh, Amit Sharma, Samrat Mondol

## Abstract

Accurate assessment of demographic parameters is crucial for conserving endangered species. The Greater One-horned Rhinoceros (*Rhinoceros unicornis*), a sexually monomorphic species, lacks a validated molecular sexing protocol for diverse biological samples. This study established a multi-marker molecular sexing protocol using blood, tissue, horn, and dung samples from all wild populations in India. The optimized multiplex strategy that involves three sex-linked markers: ZF, AMESC, and TSRY, enabled simultaneous amplification of X-and Y-chromosome-specific fragments in degraded samples. Testing with 286 individual rhino samples from seven parks (5 blood, 180 tissue, seven horn, and 94 dung) led to a ∼92% (195 M: 69 F) success rate. Overall, this approach validates a molecular sexing protocol with increased reliability and fewer amplification failures, facilitating large-scale demographic studies. Future research should utilize this method to examine the sex ratio of all existing Indian rhino populations across their range, ensuring their future persistence and management.

## Introduction

Long-term monitoring of wild population patterns and dynamics is critical for developing effective conservation strategies (Gibbs 2000; Sastre et al. 2009; Petrou et al. 2024). Among the numerous measurable biological variables in wild populations, demographic parameters like sex ratio assessments are essential for understanding population growth rates, parental care, kin selection, and sex-specific dispersal and mortality (Schweizer et al. 2007; Grüebler et al. 2008; Wang and Lu 2011; Lambert et al. 2021; Tyson et al. 2022). However, accurately estimating demographic parameters (sex identification, sex ratio, etc.) is often challenging for elusive (Kim et al. 2008; Stanton et al. 2015) and sexually monomorphic species (Peppin et al. 2010; Ahlering et al. 2011), and standard field observations or photography-based assessments have often proven ineffective for them (Kelly et al. 2008; Oliveira-Santos et al. 2010). Consequently, the introduction of molecular sexing approaches has emerged as a quick, inexpensive, and reliable substitute (Robertson and Gemmell 2006; Bidon et al. 2013) to address some of these challenges related to many taxa of interest (Primates-Villesen and Fredsted 2006; Morrill et al. 2008; Takabayashi and Katoh 2011; Rodents-Clapcote and Roder 2005; McFarlane et al. 2013; Leporids-Fontanesi et al. 2008; Chiroptera-Korstian et al. 2013; Zarzoso-Lacoste et al. 2018; Felids-Ortega et al. 2004; Pilgrim et al. 2005; DeCandia et al. 2016; Canids-Fernando and Melnick 2001; Sastre et al. 2009; Bovids-Pfeiffer and Brenig 2005; Gokulakrishnan et al. 2012, 2013; Cetaceans-Morin et al. 2005).

Traditionally, various approaches have been employed for reliable molecular sexing in mammals (XX/XY system). The most widely used methods include (a) amplification of Y-linked markers with an external control (in the form of nuclear-(Hasegawa et al. 2000; Pilgrim et al. 2005; Lamb et al. 2014; DeCandia et al. 2016), mitochondrial-(Taberlet et al. 1993; Gupta et al. 2006; Joshi et al. 2019), or protein markers- (Vozdova et al. 2019)); (b) single or multiple XY-linked marker amplification (Sullivan et al. 1993; Pande and Totey 1998; Fernando and Melnick 2001; Vidya et al. 2003; Shaw et al. 2003; Peppin et al. 2010; Stoops et al. 2018). These approaches mitigate significant concerns related to false negatives and have proven advantageous for non-invasive samples from free-ranging, elusive animals (Bidon et al. 2013; Pelizzon et al. 2017). Despite this, there is still a lack of species-specific protocols for many endangered species worldwide.

In this study, we developed a novel, multi-marker-based quick, and robust molecular sexing approach for the Greater One-horned Rhinoceros (*Rhinoceros unicornis*, hereafter called the Indian rhinoceros). The Indian rhinoceros is an example of a typical sexually monomorphic species that has experienced a complex population history across the Indian subcontinent (Ghosh et al. 2022). The wild populations (∼4000 individuals) currently reside across fragmented protected habitats in India (seven parks covering the states of Assam, West Bengal, and Uttar Pradesh) and Nepal (four parks namely Chitwan National Park (NP), Bardia NP, Suklaphanta NP and Parsa NP) (Talukdar 2024). Significant conservation and management efforts have led to population recovery across the species’ range, and the subsequent efforts focus on population management through translocation initiatives (Melletti et al. 2025). Information on the sex ratio would be critical in this regard, but no systematic data is available from their distribution due to logistic difficulties in sex identification. Available molecular sexing approaches (Peppin et al. 2010; Borthakur et al. 2016; Stoops et al. 2018) do not adequately address the concerns of misidentification from poor-quality samples. We addressed this gap by presenting a robust, reliable sexing method that can be used for diverse types of biological samples collected at the population level. We believe that this approach can be successfully used in assessing sex ratios, movement patterns, and other biological parameters that need demographic details.

## Materials and methods

### Permission and ethical considerations

The Ministry of Environment, Forests and Climate Change (MoEF&CC), Government of India, granted all permissions for tissue sampling under the RhoDIS-India program (Letter No. 4-22/2015/WL). Permissions for dung sampling from all rhino-bearing parks have been issued by respective state forest departments (Assam: A/GWL/RhoDIS/2017/913, 3653/WL/2W-525/2018, WL/FE.15/22; West Bengal: 3967/WI/2W-525/2018; and Uttar Pradesh: 1978/23-2-12 (G). Ethical permissions for tissues were not necessary as they were collected from naturally deceased rhinos by the respective state forest departments during post-mortem or management procedures.

### Sample collection, DNA extraction, and individual identification

Different types of rhino biological samples (blood, tissue, horn, and dung) were used in this study to examine the effects of DNA quality during molecular sexing. A total of 311 samples (blood-5, tissue-180, horn-7, and dung-119, respectively) were collected, representing all extant rhino populations across their Indian distribution. DNA from 133 individual rhinos (∼47% of the total sample size, 5 blood, 119 tissues, 4 horns, and 5 dung samples, respectively) was already available to us (See (Ghosh et al. 2021, 2022)), while 178 new rhino samples (61 tissue, 3 horn, and 114 dung samples) were added from different rhino-bearing parks of Assam, West Bengal, and Uttar Pradesh. For all newly collected rhino samples, DNA was extracted in the laboratory following established protocols (blood, tissue, and horn - Ghosh et al. 2021; dung -Biswas et al. 2019). For tissue and horns, finely macerated samples were lysed overnight with 300μl ATL buffer and 30μl proteinase K (20mg/ml) at 56°C. During the horn lysis, 30μl of DTT (1M) solution was added. Dung DNA was extracted from the swabs, which were incubated overnight at 56°C with 700μl ATL buffer and 60μl (20mg/ml) proteinase K for lysis. For all sample types, the subsequent steps were performed as per QIAamp DNA Tissue Kit (QIAGEN Inc., Hilden, Germany) protocols. DNA was eluted twice (100μl volume each) using TE buffer (pH 7.8) and stored at -20°C. An extraction negative was included with each extraction set to monitor possible contamination. All dung samples were processed in a physically separate, non-invasive, sample-dedicated laboratory space to reduce any chances of contamination. Before further processing, all extracted blood, tissue, and horn DNA samples were quantified using a spectrophotometer (Promega Corporation-QuantiFluor® ONE dsDNA System, USA).

For individual identification, the established protocol for the Indian rhino that employs a 12 microsatellite loci panel (Ghosh et al. 2024) was utilized. PCRs were performed in reaction volumes of 10 μl containing 4 μl of 2X Qiagen multiplex PCR buffer mix (QIAGEN Inc., Hilden, Germany), 0.2 μM labeled forward primer, 0.2 μM unlabeled reverse primer, 4 μM BSA (4 mg/mL), and 2 μl of rhino DNA at a concentration of 10 ng/μl. The previously selected ‘genotyping reference sample’ for the RhoDIS-India program (Ghosh et al. 2021) was used as a positive sample to ensure data call uniformity for individual identification. PCR and extraction negatives were included in each reaction set to monitor for contamination. Amplified PCR products were visualized in 2% agarose gels and further genotyped using LIZ 500 size standard (Applied Biosystems, California, USA) in an ABI 3500XL Genetic Analyzer (Applied Biosystems, California, USA) and were scored manually using GENEMARKER (Softgenetics Inc., Pennsylvania, USA) with already established allele bins (Ghosh et al. 2021).

### Marker selection for molecular sexing

Through a comprehensive literature survey, we selected five potential sexing markers (Hasegawa et al. 2000; Peppin et al. 2010; Borthakur et al. 2016; Stoops et al. 2018; Lim et al. 2020) (Table 1) for extensive testing on Indian rhinos. The shortlisting was based on the following criteria: (i) markers were used in rhino or closely related species or taxa, (ii) amplified shorter fragments (<250 bp) for higher amplification success in degraded samples (Taberlet et al. 1999), and (iii) targeted both X and Y chromosome regions to reduce false-negative errors (Pagès et al. 2009). All initial PCR standardizations were conducted using a set of 14 reference rhino samples with known sex (7M:7F). The primers were standardized using various combinations (n=6; AMEL & ZF; AMESC & ZF; AMEL & RSRY & TSRY; AMESC & RSRY & TSRY; ZF & TSRY & RSRY; and all five primers together) to ensure the most unambiguous sexing results. PCRs were performed in 10 μl reactions for each primer, containing 4 μl of Qiagen Multiplex PCR Master Mix (QIAGEN Inc., Hilden, Germany), 0.25 μM of primer mix, 3 μl of BSA (4 mg/ml), and 2 μl of rhino DNA (1:10 diluted). PCR conditions included an initial denaturation (95°C for 15 min); 40 cycles of denaturation (95°C for 30 s), annealing (45-60°C gradients for 40 s), and extension (72°C for 40 s); followed by a final extension (72°C for 10 mins). PCR and extraction negatives were included to monitor contamination. The amplified products were visualized on a 3% agarose gel, and any primer producing no or non-specific amplifications was discarded.

**Table 1.**
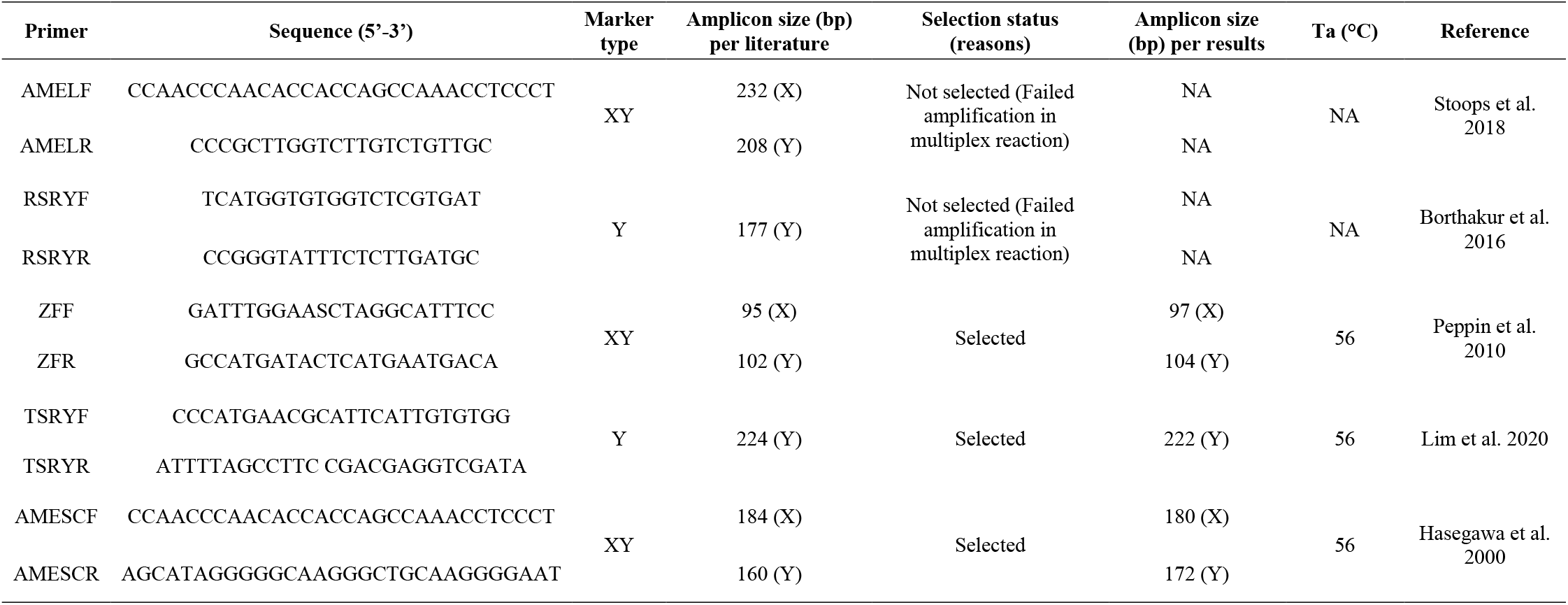
Molecular sexing markers found in literature for the Indian rhinoceros.

Due to the similar amplicon sizes found in some primers, we employed both gel and capillary-based methods during the initial standardization. The results generated from all primer combinations were repeated independently three times and compared to determine consensus results. For the capillary-based sexing approach, marker-specific allele bins were created for uniform and unambiguous calling. Finally, the standardized multiplex sexing panel was used to generate data from field-collected rhino samples using the previously mentioned multi-tube approach.

## Results

Out of the 178 field-collected samples tested in this study, we identified 153 unique individuals (61 tissue, three horn, and 89 dung samples). Combined with the previously available 133 individuals (Ghosh et al. 2021, 2022), we used 286 individual rhino samples (blood-5, tissue-180, horn-7, and dung-94) for molecular sexing.

Initial standardization with five shortlisted primers yielded the expected banding patterns in reference samples (n=14; 7M:7F). Among the six tested multiple combinations (AMEL & ZF; AMESC & ZF; AMEL & RSRY & TSRY; AMESC & RSRY & TSRY; ZF & TSRY & RSRY; and all five primers together), the capillary-based genotyping approach produced clear and consistent profiles in three-primer combinations (ZF, AMESC & TSRY), which were selected for further analyses. This multiplex assay enabled simultaneous detection of two X-linked and three Y-linked markers (Fig. 1) (ZF: X-97 bp, Y-104 bp; AMESC: X-180 bp, Y-172 bp; TSRY: 222 bp). Applying the optimized multiplex assay to 286 samples led to successful sex identification for 264 samples (∼92.3% success rate), including 195 males and 69 females. The remaining 22 samples (∼7.7%, all field-collected dung samples) failed to amplify. Sample type-wise success rates for blood, tissue, horn, and dung were 100%, 93.3%, 100%, and 89.3%, respectively. Among the markers, ZF showed the highest amplification success (98%), followed by AMESC (89%) and TSRY (73%).

**Figure 1.**
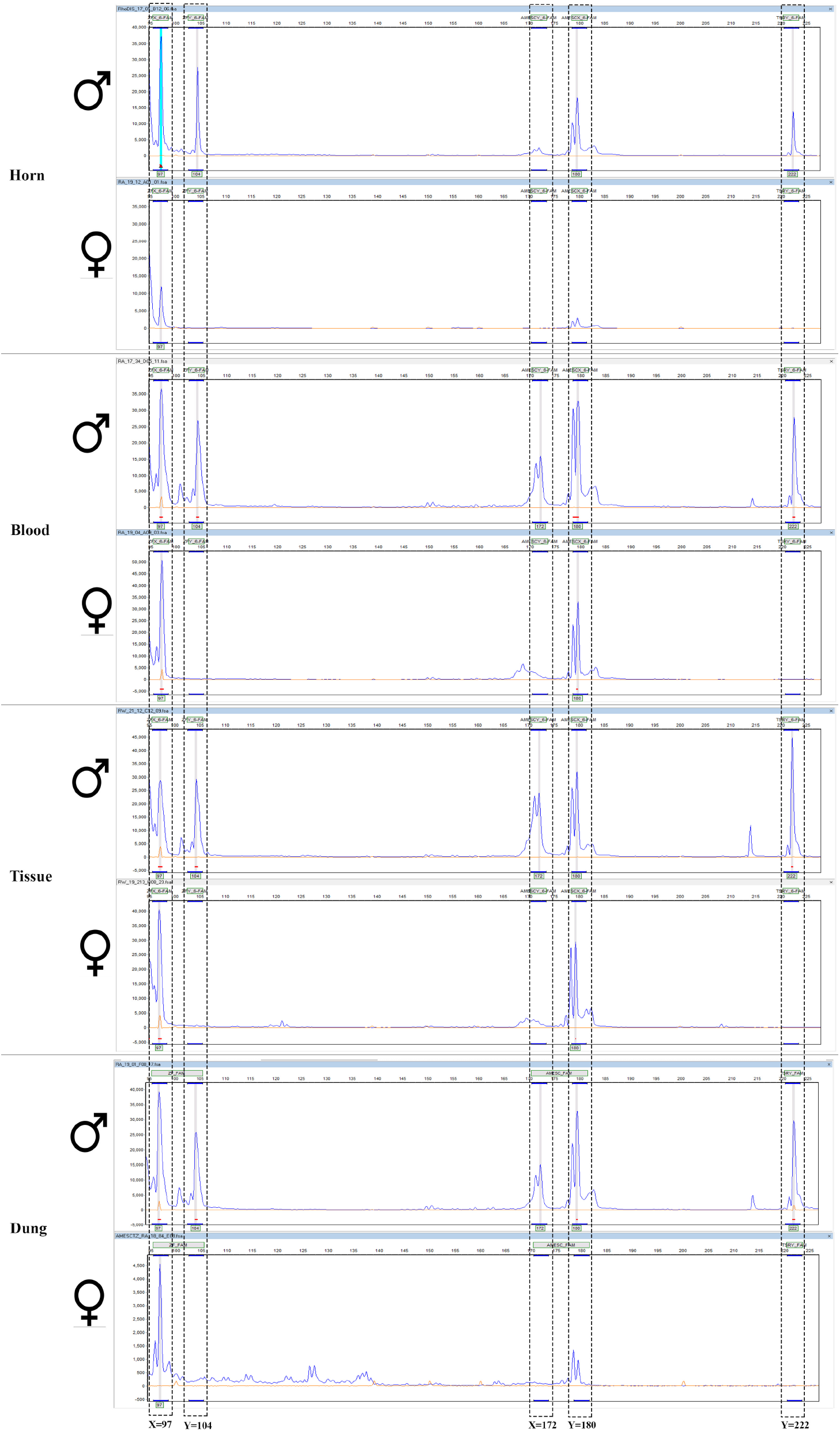
Representative electropherograms of molecular sexing outputs from male and female rhinos. Results from different types of biological samples (horn, blood, tissue, and dung) are shown here.

## Discussion

Standardizing a multiplex PCR protocol for sex determination enables efficient large-scale data generation, crucial for understanding species’ ecological and demographic processes (Robertson and Gemmell 2006). The molecular sexing method also helps assign sex to forensic or historical samples collected without sex information (Bidon et al. 2013). Assessing the precision of sex identification is generally a two-phase approach (Robertson and Gemmell 2006), involving evaluation of the sex-specificity of the test on a few reference samples and determining the test’s accuracy in a greater number of samples (Lessells and Mateman 1998; Griffiths 2000). In this regard, our study demonstrates a validated molecular sexing approach for the Indian rhinoceros, yielding reliable and reproducible results from various sample types (tissue, blood, horn, and dung). During the standardization process, we observed successful amplifications with AMEL and RSRY markers individually, but they consistently failed in multiplex reactions and were therefore discarded. Our final multiplex marker panel, which includes short amplicon sizes, concurrently amplifies both X- and Y-linked fragments. This methodology enhances amplification from compromised DNA samples and aids in internal cross-verification, thereby diminishing the probability of false negative errors (Taberlet et al. 1999; Zenke et al. 2022). The overall success rate of ∼92.3% underscores the robustness of the marker panel developed in this study. While tissue and blood samples showed near-complete success, the failure in a subset of dung samples (7.7% of the total dataset) may be attributed to DNA degradation, suboptimal DNA concentrations, or the presence of PCR inhibitors (Taberlet et al. 1999; Reddy et al. 2012). We observed a direct relationship between the marker amplicon sizes and amplification success (ZF - 98% success rate, AMESC - 89%, and TSRY - 73%, respectively), as anticipated.

We also evaluated two-marker combinations instead of the full three-marker panel to offer more choices to researchers with limited resources. The following amplification success rates were observed for sex identification in tissue and dung samples, respectively: AMESC-TSRY: 63.54% and 28.72%, AMESC-ZF: 56.77% and 30.85%, and ZF-TSRY: 92.19% and 92.55%. Although two-marker combinations can be a cost-effective alternative, we recommend the three-marker system because of its greater reliability in minimizing false-negative errors, particularly when dealing with low-quality samples. However, it is important to note that the three-marker multiplex assay would require a capillary-based genotyping approach for the best results. These markers exhibited slightly different amplicon sizes compared to previously published reports on Indian rhinos (Peppin et al. 2010- n=2; Borthakur et al. 2016- n=10; Stoops et al. 2018- n=8), possibly due to the limited sample sizes used in those studies. We recommend that future studies use large and diverse sample sets for standardization of any such approach (Waits and Paetkau 2005).

Genetic tools, especially when combined with non-invasive sampling, offer valuable insights for conservation and management (Waits and Paetkau 2005). Despite advances in molecular sexing for endangered species, reliable methods are still lacking for many species globally. Recent works highlight the use of molecular tools in rhino biology (species evolution, individual identification, population structure, etc.), and the inclusion of accurate demographic information (developed in this study) will uncover detailed population-specific patterns across their range. Based on the results of this pilot study, future research should expand to apply this approach to all existing rhino populations across India for their future management. For a long-lived, slow-reproducing species like the Indian rhino, such synergistic efforts may be the key to its future persistence.

## Acknowledgements

We thank the Ministry of Environment, Forest and Climate Change (MoEF&CC), Government of India, for their invaluable support in implementing this project. Thanks are also extended to the Forest Departments of Assam, West Bengal, and Uttar Pradesh for providing necessary permits and help while collecting the field samples and for providing the tissue samples. We thank the World Wide Fund for Nature-India (WWF-India) team for their financial, field and logistic support. Our appreciation goes to Ankit Pacha, Dr. Shrutarshi Paul, and Dr. Shiv Kumari Patel for their expert opinions during primer standardization. We thank Dr. C.P. Sharma for his assistance with reference rhino horn sampling. Finally, we thank the Director, Dean, Research Coordinator, and Nodal Officer of the Wildlife Forensics and Conservation Genetics Cell for their support.

## Author contributions

SM designed the study, while SM and AS secured the necessary funds and obtained permissions to conduct the study. Sample collection was carried out by TG and SK, with supervision from AS. SK contributed to data curation, generation, and analysis. SM, SK, and TG performed data visualization. Both SK and SM wrote the manuscript with contributions from TG and AS. All authors read and approved the final manuscript.

